# Polymer physics of structural evolution in synthetic yeast chromosomes

**DOI:** 10.1101/2022.09.14.507906

**Authors:** Giovanni Stracquadanio, Kun Yang, Jef D. Boeke, Romain Koszul, Joel S. Bader

## Abstract

Yeast SCRaMbLE is an experimental method using designed synthetic yeast chromosomes to generate combinatorial diversity through genome rearrangements. These events occur at designed loxPsym recombination sites through the activity of Cre recombinase. While the synthetic SCRaMbLE system was designed to explore minimal genomes and permit rapid genome evolution, the pattern of recombinations also reflects inherent properties of DNA looping required to coalesce pairs of loxPsym sites.

Genomes of yeast strains generated by SCRaMbLE are analyzed here using a new statistical mechanics model, called the SCRaMbLE Polymer Interaction (SPI) model. SPI uses polymer physics to model recombinations, and implements efficient rejection sampling and histogram reweighting algorithms to conduct SCRaMbLE experiments *in silico*.

Using SPI, we found that recombination events observed experimentally are consistent with a random walk scaling exponent ranging between 0.45 and 0.6, which spans values of 0.5 for a Gaussian polymer and 0.588 for a self-avoiding walk. SPI provides a highly accurate tool to study SCRaMbLE recombinations and massively parallel genome recombination experiments.

## Introduction

A fundamental problem in biology is the identification and characterization of the minimal genetic information required for cellular life [1, 2, 3]. While evolutionary arguments indicate that strictly non-essential genes will be lost, under favorable conditions, with unlimited rich nutrients and absent stresses, genomes may be minimized. For example, bacterial intracellular parasites and endosymbionts, with highly controlled intracellular environments, have evolved very small genomes through gene loss [4, 5, 6, 7, 8]. The parasitic bacterium *Mycoplasma genitalium*, which lives in epithelial cells, has the smallest known genome among organisms that can grow independently in axenic culture [9]; this bacterium’s 580 Kb genome contains only 482 protein-coding genes [10]. Symbiotic bacteria require even less genetic information for life [11, 12]; for instance, the *Candidatus Tremblaya princeps*, an endosymbiont of the citrus mealybug *Planococcus citri*, has a 139 Kb genome with just 121 genes [13].

These studies have consistently shown that genes involved in DNA replication, transcription, and protein synthesis are largely conserved across different species, while genes responsible for organism-specific metabolism can vary. Based on this principle, comparative genomics approaches have proposed minimal gene sets required for life [14]. However, these statistical methods are limited by the number of taxa available for inference and the quality of gene function annotation.

Conversely, large-scale knock-out studies have been performed in many organisms with the goal of identifying genes that, when deleted, do not lead to a viable cell; this information is used to identify set of genes that are individually essential for cellular life. For example, in *Saccharomyces cerevisiae*, the best-characterized eukaryotic organism, about 17% of the 6, 000 annotated protein-coding genes are essential for growth in rich media [15]. Although the remaining genes can be deleted individually, simultaneous deletions can be lethal due to the accumulation of fitness defect or genetic interactions. While synthesis and testing of all possible pairwise deletion combinations would therefore be highly informative, this brute force approach already approaches the limits of experimental feasibility for a single condition and is not feasible for higher order combinations.

Harnessing recent advances in synthetic biology and DNA synthesis, the *Saccharomyces cerevisiae 2*.*0* (Sc2.0) project aims to identify a minimal eukaryotic genome using a redesigned yeast genome. Specifically, an *in vivo* evolutionary system named Synthetic Chromosome Recombination and Modification by loxP-mediated Evolution system (SCRaMbLE) is embedded in the genome of synthetic yeast [16, 17, 18]. Each non-essential gene of the synthetic genome carries at its 3’ end (just downstream of the stop codon) a loxPsym site, a synthetic 34 bp palindromic DNA sequence. Upon expression of Cre recombinase, pairs of loxPsym sites can interact essentially at random, leading to inversion or deletion of the intervening genomic region, or more complex rearrangements. The genome rearrangements generated by SCRaMbLE provide diversity that can be used to select for desired phenotypes. Phenotypes corresponding to minimal genomes include gene content and chromosome length; in other contexts, fitness may correspond to the ability to survive in an extreme environment or to generate a desired metabolite and in these cases duplication of genes via SCRaMbLE can lead to a “gain of function”.

Sequencing strains subjected to SCRaMbLE under selection provides information that can be used to identify the most important genes or rearrangements. However, performing this analysis requires a model for the recombination events sampled by SCRaMbLE, which are random in principle but may be constrained physically by factors such as DNA looping and stiffness, tethering of chromosomes to protein complexes, membranes or other DNA molecules. The loxPsym sites participating in a recombination events are brought into contact by DNA looping, and the physical constraints of looping introduce a distance-based recombination probability. Recombination hotspots and coldspots may exist as well, as they do for meiotic crossing over; identifying and distinguishing these deviations from random recombination events biased by phenotypic selection requires development of a null model for neutral SCRaMbLE recombination events that do not affect cellular fitness. Such physics-based model could have general value in predicting probabilities of DNA looping structures relevant to enhancer/promoter interactions, chromatin structure, and chromosome conformation capture.

Here we introduce the SCRaMbLE Polymer Interaction (SPI) model to study synthetic genomes generated by SCRaMbLE. Our model adapts the contact probability for polymers used for DNA looping to estimate the probability of contacts between pairs of loxPsym sites [19, 20]. Parameters in the model correspond to the long-range scaling exponent for polymer mean square distance, *ν*, the shortrange persistence length of DNA, *b*, and the mean number of recombination events per genome, *λ*. To estimate model parameters, we developed a rejection sampling algorithm and an efficient histogram reweighting method to enable large scale parameters exploration [21].

We then tested and validated SPI using whole-genome sequencing data of 64 synthetic yeast strains containing a circular version of the synthetic right arm of chromosome IX, *synIXR*, which have been previously rearranged using SCRaMbLE [22]. We examined the concordance between the rearrangements in the experimental strains and in strains simulated over a range of parameter values. We particularly explored values of the scaling-exponent *ν* ranging from 0 (uniform probability) to 0.5 (non-self-avoiding walk) to 0.588 (numerical estimate for the self-avoiding-walk in three dimensions), and the persistence length parameter, ranging from 50bp to 300bp, including the widely accepted value of 150bp. After finding the best parameter settings by maximum likelihood, we compared the structure and biological features of the simulated and SCRaMbLE genomes to infer fitness requirements and minimal chromosome length. Our results confirm that SPI is an accurate model for analyzing SCRaMbLE experiments, and a useful exploratory tool to investigate minimal chromosomes conformations using only sequence information.

## Methods

### Genome structure and notation

The synthetic yeast genome is designed to generate structural rearrangements of genomic loci, also called segments, flanked by loxPsym recombination sites. LoxPsym sites were designed to occur primarily at the boundary between the coding and 3^*1*^ untranslated region of each non-essential gene, yielding segments of varying lengths.

Here we considered the circular synthetic chromosome *synIXR*, which contains *n* = 43 segments, numbered *i* = 1, 2, 3, …, *n* (see Supplementary Table 1). The structure of a rearranged genome is represented by a list of integers *G* = *{s*_*i*_*}*, where inverted segments are negative, deleted segments do not occur, and duplications and higher order amplifications yield multiple appearances of a signed segment number. Integers associated with segments containing essential genes will always be present [23]. Importantly, since the model parameters will be estimated using sequencing data for *synIXR*, which is a circular chromosome, any circular permutation or complete reversal with sign change is equivalent.

### Distance-dependent recombination probability between loxPsym sites

Although deletions and inversions should occur with equal probability, an assumption already exploited by other models [24], the number of segments involved in a recombination event depends on the probability of interaction between two loxPsym sites. Here we represent DNA as a polymer of physical length *L*, effective monomer length *m*, and persistence length *b* [25, 19]. For a linear polymer, when *L > m*, the probability density *ρ*(**r**; *L*) for end-to-end vector **r** is

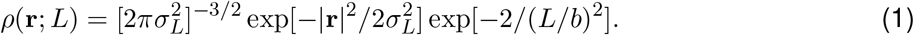

We assume that the variance follows the scaling law,

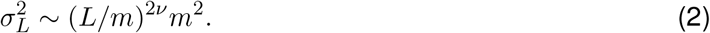

For a long polymer with *L ≫ m*, the mean square displacement is 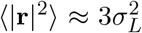. In the ideal case of a non-self-avoiding walk, the scaling exponent *ν* = 0.5. For a self-avoiding walk in three dimensions there are no exact results, but theoretical approximations and simulations both give *ν ≈* 0.588 [19]. Here we consider *ν* an adjustable parameter that can be estimated from the data.

The end-to-end contact probability from Eq. 1 for a linear polymer of length *L* is *ρ*(**0**; *L*). The contact probability *p*(*L, L*^*1*^) between two sites a distance *L* apart in a circular polymer of total length *L* + *L*^*1*^ is approximated using the linear probability density for the two regions of length *L* and *L*^*1*^ and neglecting the interactions between them,

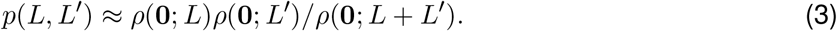

The denominator is a normalization factor that may be approximated as ∫ *d***r***ρ*(**r**; *L*)*ρ*(**r**; *L*^*1*^) and which cancels out of the final results.

For a synthetic yeast chromosome, recombination events are restricted to positions corresponding to pairs of loxPsym sites. For a pair *ij* of loxPsyms sites separated by *N*_*ij*_ nucleotides in one direction and 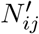 nucleotides in the other direction around the chromosome, the rate of contacts is proportional to 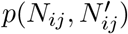 (see Eq. 3). Thus, the probability that a pair *ij* of loxPsyms sites is involved in a recombination event is the normalized probability *P*_*ij*_,

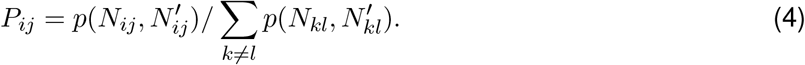

Since all powers of *m* cancel, the recombination probabilities is independent of the monomer length and symmetric in *N* and *N* ^*1*^, calculated as *P*_*ij*_ = *Q*_*ij*_*/ Σ*_*k≠l*_ *Q*_*ij*_ and

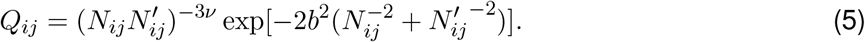

Nonetheless, the recombination probabilities do depend on *ν* and *b* (see Figure 1). While the literature value *b* = 150 bp is generally assumed for double stranded DNA (dsDNA) [26, 27], we treated *b* as a parameter to be estimated from the data. While the model above is motivated in terms of recombination probabilities for self-avoiding walks and non-self-avoiding walks, it can also describe recombination events happening uniformly at random (*ν* = *b* = 0) and other biologically interesting alternatives. This model therefore provides a unified framework for testing alternative hypotheses for DNA looping in SCRaMbLE.

**Figure 1:**
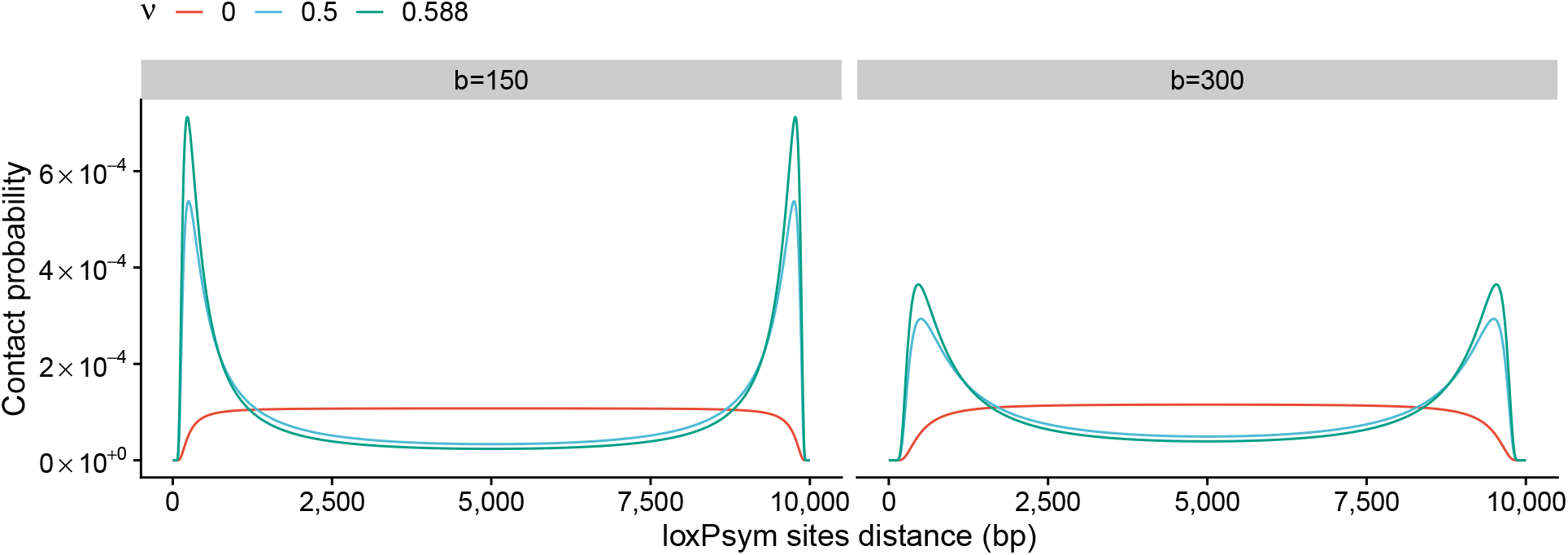
Distance-dependent contact probability between recombination sites on a circular chromosome. We report the contact probability for two recombination sites, under a random (*ν* = 0), non self-avoiding walk (*ν* = 0.5) and self-avoiding walk interaction model (*ν* = 0.588), assuming a persistence length *b* = {150, 300} bp, where 150bp represents the estimated persistence length of DNA.

### Rejection sampling and histogram reweighting

We designed a rejection sampling algorithm to generate an ensemble of chromosomes consistent with experimental constraints and phenotype selection strategies, in order to estimate model parameters and study different genomic features.

In practice, our sampling algorithm builds an initial chromosome with *n* ordered segments, which undergoes *s* recombination events with *s ∼* Poisson(*λ*). For each recombination, the pair of loxPsym sites involved was selected according to Eq. 4, and the intervening segments undergo a inversion or deletion with equal probability, as anticipated from the presence of the symmetric loxPsym site [22].

After *s* recombinations, a chromosome is accepted if it retained all the essential genes and lost at least one auxotrophy marker, mirroring the selection that was applied to the SCRaMbLE strains.

Generating large ensemble of genomes by rejection sampling is computationally expensive, which hinders large-scale parameter exploration to obtain accurate estimates. To overcome this problem, we developed a histogram reweighting method [21], which allow us to collect statistics for multiple parameters settings from a rejection sampling trajectory run with a single parameter set.

Let *F* be a property of a genome *g, F* (*g*). For a particular parameter set *θ*, the probability distribution of genomes is denoted *P*_*θ*_(*g*), and the expectation of *F* is ⟨ *F* _*θ*_ ⟩ is Σ_*g*_ *P*_*θ*_(*g*)*F* (*g*)*/* Σ_*g*_ *P*_*θ*_(*g*). In our case, *F* is the average number of times a segment is present in an ensemble of genomes. Let *P*_*θ*_ be a probability distribution for a set of parameters *θ*^*1*^. Formally, we can use the ensemble of genomes obtained from *P*_*θ*_ to compute ⟨ *F* _*θ*_ ⟩ as

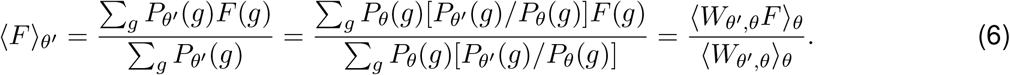

The term 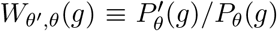 is the probability ratio for sampling a genome *g* from a distribution with parameters *θ*^*1*^ instead of *θ*. Then, it is possible to compute *W*_*θ*_^*′*^_,*θ*_ for each genome, sampling parameters *θ* = *{λ, ν, b}*, and desired parameters *θ*^*1*^ = *{λ*^*′*^, *ν*^*′*^, *b*^*′*^*}*, as

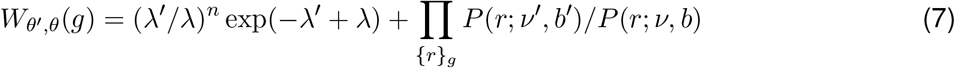

where the first term is the ratio between the expected number of recombination events, and the product over *r* is computed over the sequence of recombination events leading to the final genome structure *g*. Reweighting procedures should consider samples drawn from distributions that are close to the reweighted parameters in order to reduce statistical errors [21]. For this reason, in our experiments, for each reweighting parameter setting, we considered only genomes generated by simulations with the closest grid points in the parameter space.

For both rejection sampling and histogram reweighting, we estimated model parameters using maximum likelihood. Specifically, let *p*_*θ*_(*k*) the probability of retaining segment *k* after SCRaMbLE, as estimated by using SPI with parameters *θ*. Then, given a population *g* of SCRaMbLE strains, the log-likelihood of the model can be computed as

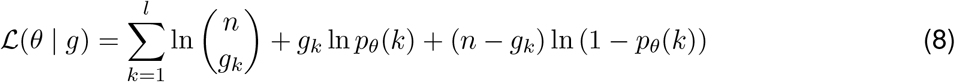

where *l* is the number of segments in the chromosome, *n* is the total number of SCRaMbLE strains, and *g*_*k*_ is the number of strains retaining segment *k*.

We then computed confidence intervals for model parameters using bootstrap. Specifically, genome replicates were sampled at random with replacements from the set of SCRaMbLE strains, and then we selected model parameters using maximum likelihood; finally, for each parameter, we report the corresponding 95% confidence interval.

## Results

We used whole-genome sequencing data of 64 synthetic yeast strains including the episomal circular *synIXR* chromosome to estimate parameters of our model. All experimental strains retained essential genes and the centromere, a feature essential for chromosome replication, and were selected to have lost at least one auxotrophic marker. Moreover, 31 of the 64 strains analyzed had at least one duplicated segment; nonetheless, since our objective is to predict minimal genomes, we did not explicitly model duplication events and counted each duplicated segment only once.

We then determined model parameters by maximum likelihood using grid search; specifically, parameter were explored on a grid, Ω = ***λ*** × ***ν*** × ***b***, where ***λ*** = [4, …, 13], ***ν*** = [0.3, …, 0.7], ***ν*** = [50, …, 300] with a step size of 1, 0.05, and 50bp respectively (see Supplementary Table 2). With this experimental design, we obtained a total of 540 parameters settings for which we generated 10^4^ genomes each.

Here we found the best model parameters to be *λ* = 9, *ν* = 0.55, *b* = 200. (log-likelihood = *−*178.713; see Figure 2A and Supplementary Table 3), with generally high-confidence (see Figure 2B). Specifically, the 95% confidence intervals for the scaling exponent *ν*, ranging from 0.45 and 0.6, suggests that recombination events induced by SCRaMbLE follow a self-avoiding walk but not a uniform random process. The persistence length *b* instead is bound to 200 bp, suggesting recombination event probability scales proportionally to the expected persistence length of DNA.

**Figure 2:**
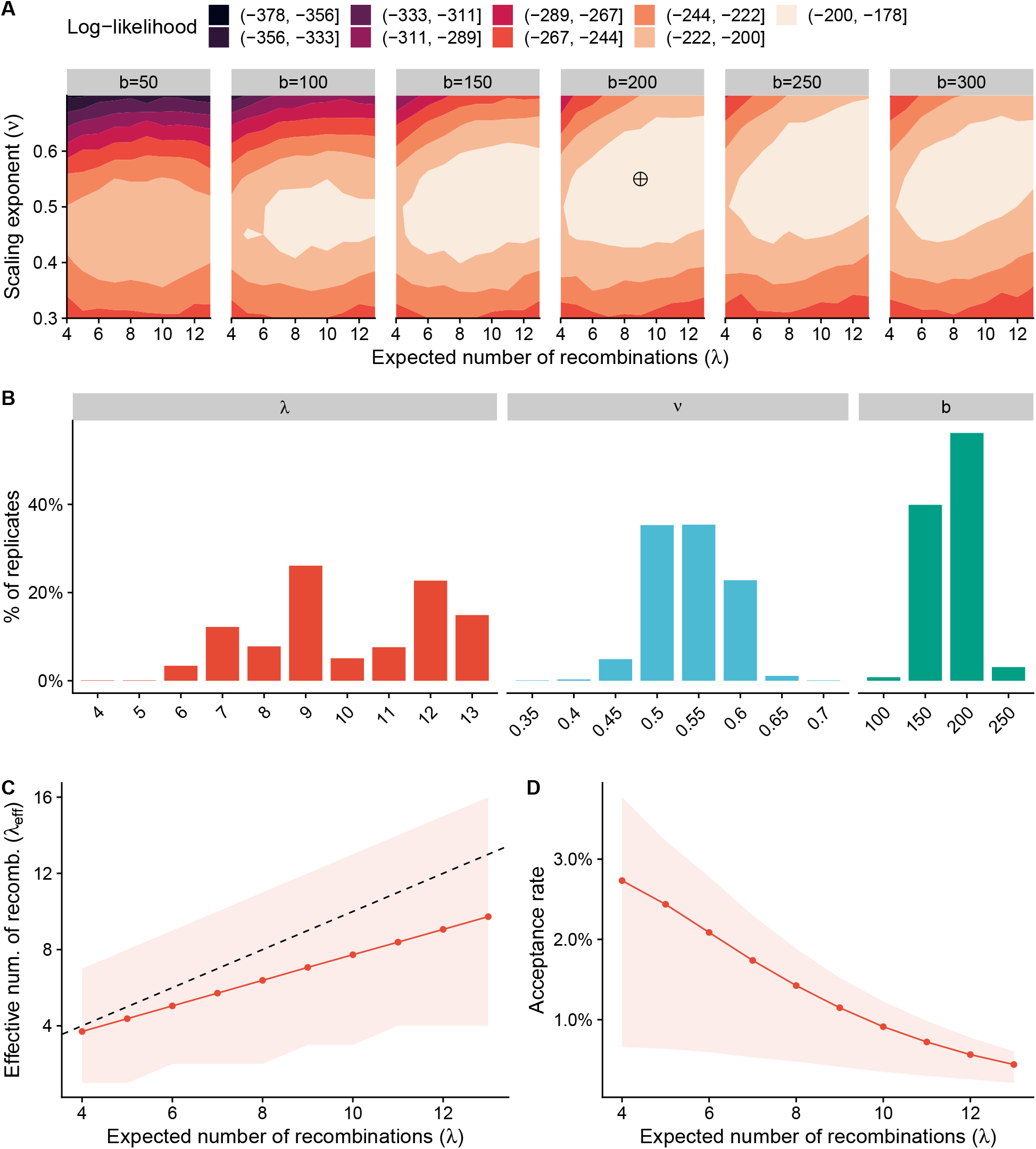
Model parameter estimation and analysis using rejection sampling. **A)** We report the log-likelihood value for each parameter setting, while denoting with a crossed circle the best parameter setting.**B)** bootstrap confidence intervals for each model parameter. **C)** We compare the average expected and the effective number of recombination events for various *λ* values; the area in light red represents the 95% confidence interval of the effective number of recombination events. **D)** Analysis of the percentage of genomes accepted during sampling at various *λ* values; the area in light red represents the 95% confidence interval of the number of percentage of accepted genomes.

The 95% confidence region for the recombination Poisson parameter was broad, with *λ* ranging from 6 to 13 events. While this result supports an iterative recombination process induced by SCRaMbLE, the estimates appear unrealistic as they would cause an excessive number of chromosomal aberrations unlikely to be compatible with life. Therefore, we examined the effective number of events, *λ*_eff_, that is the number of recombination events observed in a viable genome (see Figure 2C).

Here we found that the effective number of recombination events to be *λ*_eff_ ≤ 10, fewer than the expected number of events for a Poisson random variable. This phenomenon is also confirmed when looking at the percent of accepted genomes during our sampling process (see Figure 2D), which is 2-fold higher when *λ* ≤ 10 compared to any higher setting, consistent with the downward shift of *λ*_eff_. Taken together, this result suggests a potential upper-bound on the number of recombination events.

Successively, we tested whether explicitly modelling the interaction between loxPsym sites with a polymer physics model provides more realistic genome ensembles compared to a null model, where recombination sites can interact uniformly at random. A null model is equivalent to SPI for *ν, b →* 0. Therefore, we fixed *ν* = *b* = 0, and we fit a null model using maximum likelihood to estimate the number of events *λ*, over the space ***λ*** = [3, …, 15] with a step size of 1. We found that the expected number of events for the null model to be *λ* = 12 (log-likelihood= *−*385.237, see Supplementary Figure 1). We then performed a likelihood ratio test between SPI and the null model, and we found that our model fits significantly better than the null (*χ*^2^ = 413.048, p = 2.03 × 10^*−*90^).

### Histogram reweighting enables fine-grained analysis of the scaling exponent and persistence length parameters

We then focused on obtaining accurate estimates of the scaling exponent and persistence length parameters, using our histogram reweighting method and genomes previously obtained by rejection sampling. Here we reweighted over a grid of parameters bounded by the 95% confidence intervals estimates for the *ν* and *b* parameters, using a step size of 0.01 and 10bp respectively, while keeping the number of recombination events fixed at *λ* = 9, which is the maximum likelihood estimate obtained by rejection sampling (see Supplementary Table 2). Using genomes generated by rejection sampling with the closest grid points in the parameter space, we performed reweighting of 176 parameter settings over *≈* 9 × 10^4^ genomes.

After reweighting, we found the best model parameters to be *λ* = 9, *ν* = 0.55, *b* = 200 (loglikelihood: *−*179.295; see Figure 3A and Supplementary Table 3), which are the same obtained by rejection sampling. Analysis of the bootstrap 95% confidence intervals confirms estimates obtained by rejection sampling, with the scaling exponent being close to the known estimate of a self-avoiding walk polymer, whereas we obtained higher confidence on the persistence length parameter being close to the known estimate for DNA (95% CI: [150bp, 210bp]; see Supplementary Table 3).

**Figure 3:**
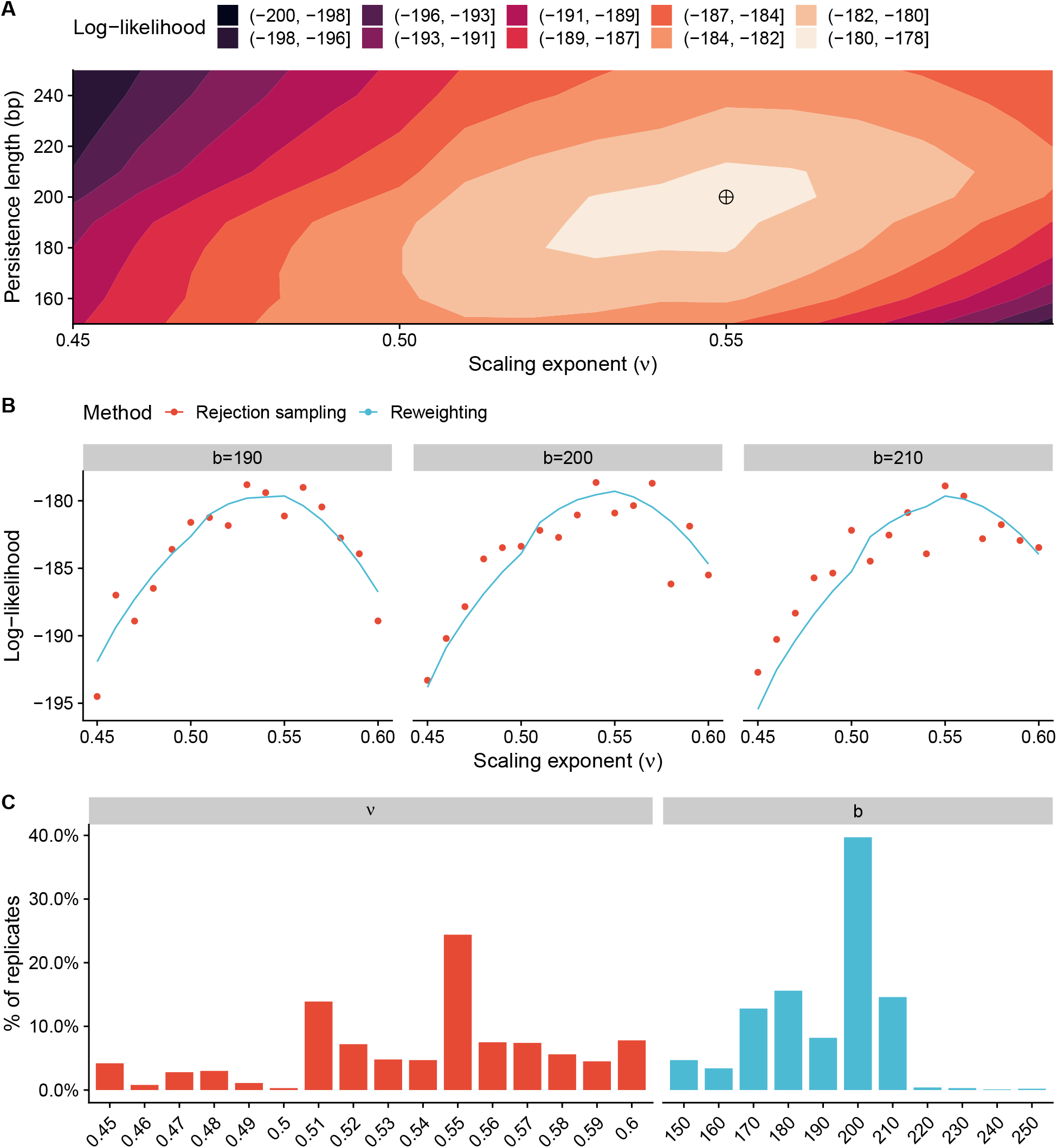
Model parameter estimation and analysis using histogram reweighting. **A)** We report the log-likelihood value for each parameter setting, while denoting with a crossed circle the best parameter setting. **B)** Analysis of log-likelihood estimates obtained by reweighting and sampling for different *ν* and *b* values at *λ* = 9, which is the maximum likelihood estimate. **C)** Bootstrap confidence intervals for each parameter.

We validated these results by performing rejection sampling over the parameters sets used for reweighting, and found the root mean squared error for the log likelihood to be 1.985 (see Figure 3B and Supplementary Figure 2), confirming that reweighting is an accurate method for extensive parameters exploration with limited computational burden (see Figure 3C).

### Discrepancies in deletion frequency can identify functional genomic regions

We then studied whether simulated genomes obtained by our model have structural and biological features similar to the population of SCRaMbLEd strains.

To do that, we first tested whether simulated genomes differed from the population of SCRaMbLE strains with respect to genome length (see Figure 4A); here we found no statistically significant difference (Wilcoxon rank sum test, W=364626, p=0.05409), with limited discrepancies observed only for extremely short genomes. Analogously, we did not find any statistically significant difference in the number of segments deleted per strain (see Figure 4B; Wilcoxon rank sum test, W = 277474, p= 0.06559), which confirms that our model generates ensembles of genomes with structural features similar to SCRaMbLE strains.

**Figure 4:**
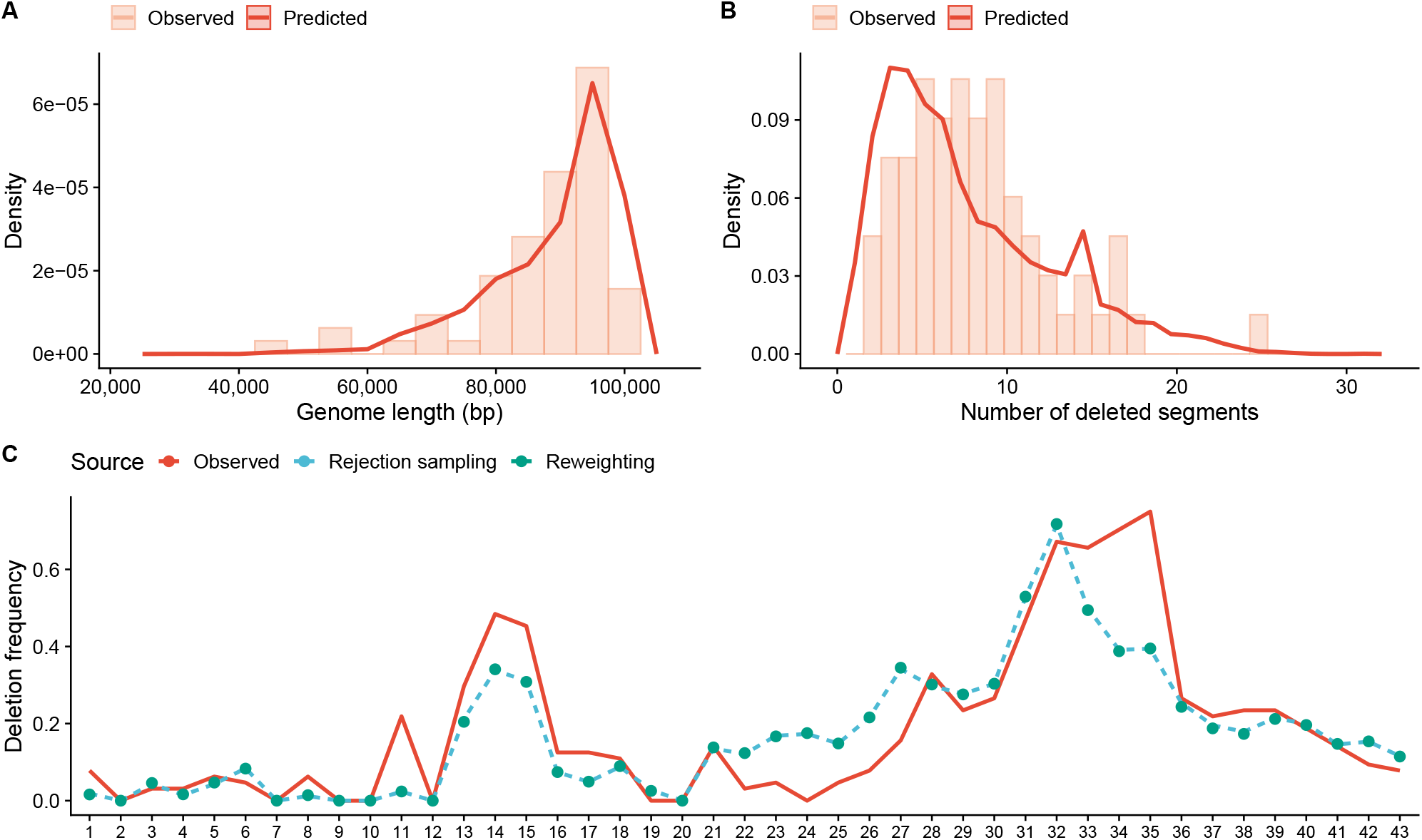
Structural and biological features analysis of simulated genomes. **A)** Analysis of genome length and **B)** segments deletion distribution in simulated and observed SCRaMbLE genomes. **C)** Deletion frequency of each segment estimated by sampling and reweighting compared to the distribution observed in SCRaMbLE genomes.

We then compared the expected frequency of deletion of each segment with the observed deletion frequency in SCRaMbLE strains, and found that our model is able to consistently describe patterns of deletions observed after a SCR_A_M_B_LE experiment, either using rejection sampling or reweighting (see Figure 4C). Deletions around the auxotrophic markers (segment 14 and 32) are enriched as observed in SCRaMbLE strains. Essential genes are always retained (segments 2, 7, 9, 10, 12, 20), and the number of segments deleted in the intervening regions is clearly constrained by their genomic location.

We observed major deviations between the expected and observed deletion frequency in the region between segments 22 and 27. In particular, segment 24 is never deleted in SCRaMbLE strains, whereas it is deleted in *≈* 20% of the simulated genomes. The discrepancy is explained by the presence of the Yeast vaccinia virus VH1 Homolog (*YVH1*) gene, the knock-out mutation of which confers a slow-growth phenotype; since our population of SCRaMbLE strains were selected to have near wild-type fitness, deletions of the *YVH1* are not represented.

Taken together, we have shown that our model simulates genomes with structural and biological features comparable to SCRaMbLE strains, and represents a an accurate model to identify physical and biological constraints associated with cellular fitness.

## Discussion

The construction of synthetic yeast chromosomes integrating an inducible evolutionary system enabled the first large-scale genome minimization experiment [22]. We proposed a new statistical mechanics model, called the SCRaMbLE Polymer Interaction (SPI) model, which can predict minimal genomes generated by a SCRaMbLE experiment. Here we hypothesized that the number of viable minimal genome is conditioned on the probability of interaction between recombination sites. We then modelled this recombination probability using polymer physics, and developed a rejection sampling algorithm and an efficient histogram reweighting method, which we validated using whole genome-data of a SCRaMbLE experiment conducted on strains integrating the synthetic right arm of chromosome IX.

We demonstrated that our model provides accurate estimates of recombination events that can lead to minimal genomes and that loxPsym sites interact following a distance-dependent self-avoiding walk process rather than a random uniform interaction process, and it is limited by persistence length of DNA. Interestingly, the predicted distribution of the chromosome length and number of deleted loci is in strong agreement with the observed distribution in SCRaMbLE strains. Moreover, simulations suggest that the SCRaMbLE genomes might have undergone multiple recombination events; this could explain why long-range recombination events are observed despite the low interaction probability between the flanking loxPsym sites. However, this hypothesis would require further *in vivo* experiments.

The ability to identify functional biological constraints just by comparing deletion frequencies represents a powerful investigation tool. Our model represents a viable solution when looking for cis-recombination events, while trans-recombination events (translocations), introduces an additional layer of complexity; while cis-recombinations probability can be described as a function of genomic distance, for recombining sites on different chromosome this is not possible. A solution would be to introduce an additional term that adjusts the probability of interaction based on consideration of the statistical probability of favored/unfavored chromosomal conformations. While trans-recombination events can be mathematically modelled, estimating the parameters of an extended model could require a significant amount of data from high-throughput genome conformation capture experiments not yet readily available.

## Contributions

G.S., J.S.B, and J.D.B conceived the study. G.S. and J.S.B. formulated and developed the model with contributions from R.K. and K.Y. G.S. implemented and tested the model. G.S. and J.S.B. wrote the manuscript with contributions from all authors.

## Acknowledgments

This work was supported by the UKRI EPSRC Fellowship (EP/V033794/1) to G.S.

## Source code availability

The source code of the SPI model and a Nextflow workflow to perform all simulations reported in our study is available at: https://github.com/stracquadaniolab/spi-nf.

