## Supplementary Materials for "Polymer physics of structural evolution in synthetic yeast chromosomes"

### Tables

| Id | Length (bp) | Type |
|---|---|---|
| 1 | 15761 | non essential |
| 2 | 689 | essential |
| 3 | 1306 | non essential |
| 4 | 5550 | non essential |
| 5 | 1629 | non essential |
| 6 | 850 | non essential |
| 7 | 5112 | essential |
| 8 | 2547 | non essential |
| 9 | 1896 | essential |
| 10 | 5186 | essential |
| 11 | 1836 | non essential |
| 12 | 1645 | essential |
| 13 | 158 | non essential |
| 14 | 1831 | aux. marker (*MET28*) |
| 15 | 245 | non essential |
| 16 | 4042 | non essential |
| 17 | 4717 | non essential |
| 18 | 945 | non essential |
| 19 | 3260 | non essential |
| 20 | 4338 | essential |
| 21 | 179 | non essential |
| 22 | 1965 | non essential |

| Id | Length (bp) | Type |
|---|---|---|
| 23 | 135 | non essential |
| 24 | 1421 | non essential |
| 25 | 4156 | non essential |
| 26 | 1497 | non essential |
| 27 | 229 | non essential |
| 28 | 1009 | non essential |
| 29 | 2049 | non essential |
| 30 | 4476 | non essential |
| 31 | 177 | non essential |
| 32 | 1445 | aux. marker(*LYS1*) |
| 33 | 1082 | non essential |
| 34 | 1586 | non essential |
| 35 | 217 | non essential |
| 36 | 2429 | non essential |
| 37 | 4102 | non essential |
| 38 | 2925 | non essential |
| 39 | 955 | non essential |
| 40 | 998 | non essential |
| 41 | 1804 | non essential |
| 42 | 985 | non essential |
| 43 | 1006 | non essential |

**Table 1: synIXR genomic structure**. For each segment in the chromosome, we report its numeric id, its length and the type, being either essential, non-essential or auxotrophic marker.

| Model parameter | Rejection sampling parameters grid (step size) | Histogram reweighting parameters grid (step size) |
|---|---|---|
| Expected number of recombination events ($\lambda$) | $[4, \cdots, 13](+1)$ | $9$ |
| Scaling exponent ($\nu$) | $[0.3, \cdots, 0.7](+0.05)$ | $[0.45, \cdots, 0.6](+0.01)$ |
| Persistence length ($b$) | $[50\text{bp}, \cdots, 300\text{bp}](+50\text{bp})$ | $[150\text{bp}, \cdots, 250\text{bp}](+10\text{bp})$ |

**Table 2: Parameters settings used for SPI simulations.**

| Model | Log-likelihood | $\lambda$ | $\nu$ | $\mathbf{b}$ |
|---|---|---|---|---|
| Rejection sampling | $-178.713$ | $9\ (6, 13)$ | $0.55\ (0.45, 0.6)$ | $200\ (150, 250)$ |
| Histogram reweigthing | $-179.295$ | $9(-)$ | $0.55\ (0.45, 0.6)$ | $200\ (150, 210)$ |
| Null (random uniform) | $-385.237$ | $12$ (-) | $0\ (-)$ | $0\ (-)$ |

**Table 3: Parameter estimation.** For each model, we report the log-likelihood value, the best estimate for each parameter, with $95\%$ confidence intervals within brackets.

### Figures

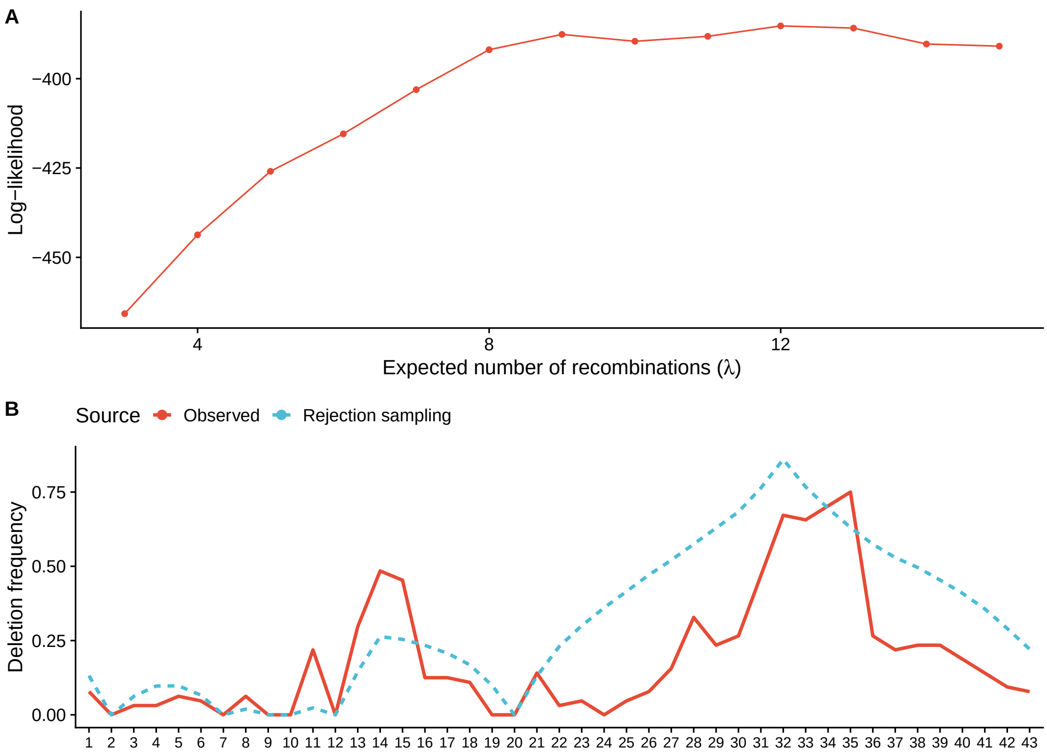

**Figure 1: Structural analysis of simulated genomes using a uniform recombination event probability model. A)** Analysis of genome length and **B)** Deletion frequency of each segment compared to the one observed in SCRaMbLE genomes.

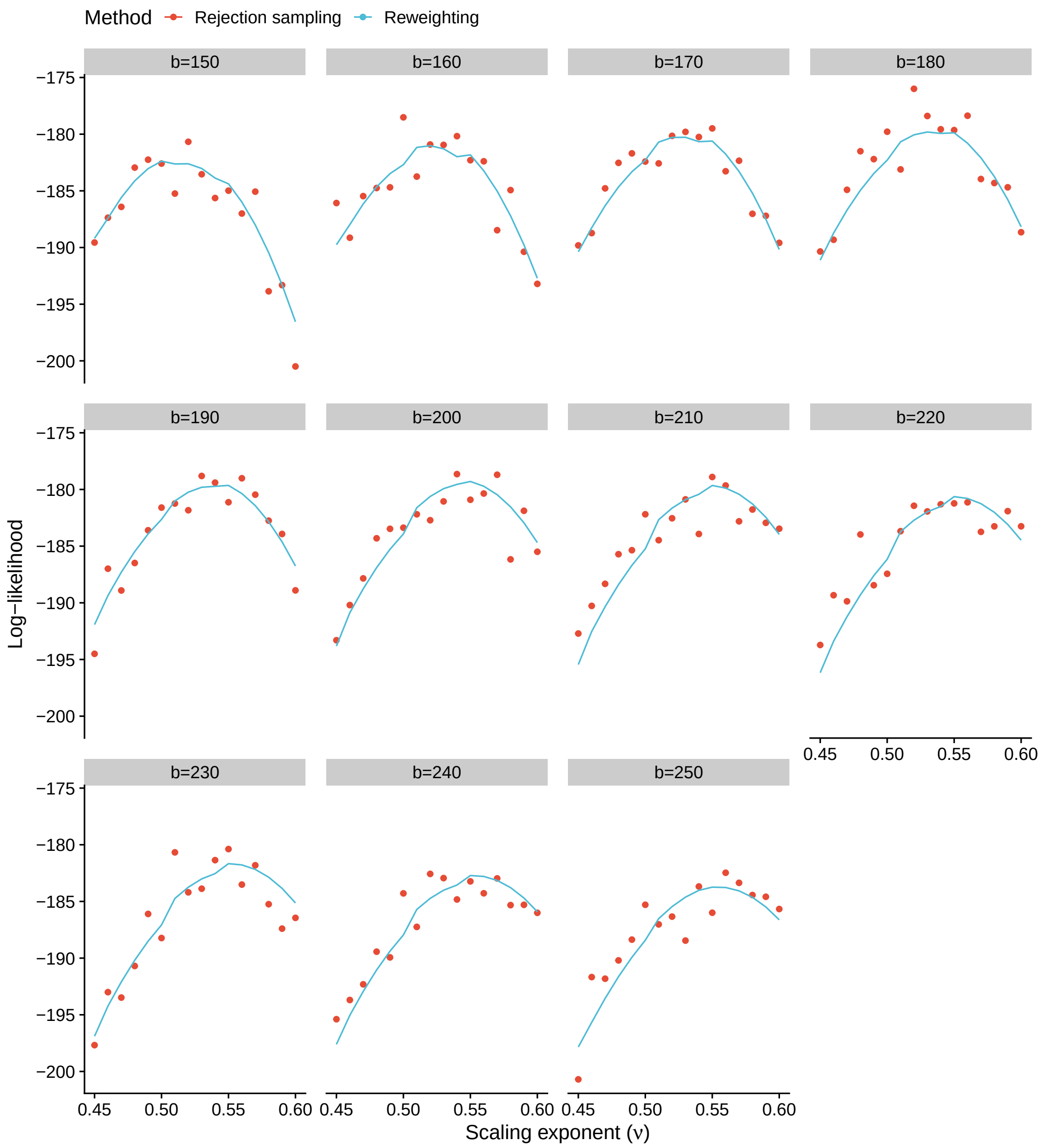

**Figure 2: Histogram reweighting performance analysis.** For each persistence length setting, we report the log-likelihood estimates of the model parameters computed by rejection sampling and histogram reweighting.
